# Self-organization of Plk4 regulates symmetry breaking in centriole duplication

**DOI:** 10.1101/313635

**Authors:** Shohei Yamamoto, Daiju Kitagawa

## Abstract

During centriole duplication, a single daughter centriole is formed near the mother centriole. The mechanism that determines a single duplication site is unknown. Here, we demonstrate that intrinsic self-organization of Plk4 underlies symmetry breaking in centriole duplication. We show that in its nonphosphorylated state, Plk4 preferentially self-assembles via a disordered linker and that this self-assembly is prevented by autophosphorylation. Consistently, the dissociation dynamics of centriolar Plk4 are controlled by autophosphorylation. We further found that autophophorylated Plk4 is localized as a single focus around the mother centriole before procentriole formation, and is subsequently targeted for STIL-HsSAS6 loading. Perturbing Plk4 self-organization affects the asymmetry of centriolar Plk4 distribution and centriole duplication. We propose that the spatial patterning of Plk4 directs a single duplication site per mother centriole.

## Introduction

The centriole is a nonmembranous organelle that is crucial for centrosome assembly and cilia/flagella formation (*1*). Centrioles are duplicated once per cell cycle and only a single daughter centriole is formed next to each preexisting mother centriole (*2*). To date, proteins that regulate centriole duplication and their protein-protein interactions have been fairly identified (*2*, *3*). Among these, Polo-like kinase 4 (Plk4) was identified as a master regulator of centriole duplication (*4*, *5*). Loss of Plk4 or inhibition of its kinase activity results in a failure to assemble procentrioles (*4*–*6*). In contrast, overexpression of Plk4 induces multiple daughter centriole formation on a single mother centriole (*7*, *8*). Plk4 phosphorylates STIL, which subsequently promotes STIL-HsSAS6 loading to the assembly site of a newly formed procentriole (*9*–*13*). In addition, Plk4 autophosphorylates its kinase domain to promote its own kinase activity (*14*) and that of its degron motif present in a flexible linker (Linker 1, L1) to enhance its own degradation via the SCF(β-Trcp) ubiquitin ligase-proteasome pathway (*15*–*18*). Despite this accumulating information, the mechanism by which the local amount and/or the kinase activity of Plk4 at mother centrioles are coordinated to ensure formation of a "single" daughter centriole on a mother centriole remains elusive.

## Results

### Autophosphorylation regulates self-assembly of Plk4 *in vitro* and in cells

To obtain new insights into the molecular basis of centriole duplication, we investigated the molecular properties of human Plk4. While performing *in vitro* biochemical experiments, we unexpectedly found that a kinase-dead (KD) mutant of a purified Plk4 fragment (Kinase + Linker 1) was highly insoluble compared with wild-type (WT) (Fig. 1A,1B and S1A). Based on this observation, we speculated that Plk4 may modulate its solubility by autophosphorylation, because Plk4 is known to autophosphorylate its kinase domain and L1 (*14*, *16*, *18*–*21*). Recent studies have indicated that protein solubility can be regulated by multiple phosphorylation within the low-complexity region (LCR) (*22*–*24*). Because human Plk4 contains an LCR in its disordered L1, we substituted some of the known and putative autophosphorylation sites in or around the LCR with alanine (10A) to generate a nonphosphorylated mutant or with glutamic acid (10E) to generate a phosphomimetic mutant, as described previously (*16*, *19*) (Fig. 1C). In line with this, the phosphorylation-induced mobility shift of Plk4 WT fragments was largely suppressed by the 10A mutation, suggesting that the target residues in WT fragments were phosphorylated *in vitro* (Fig. S1B).

**Fig. 1.**
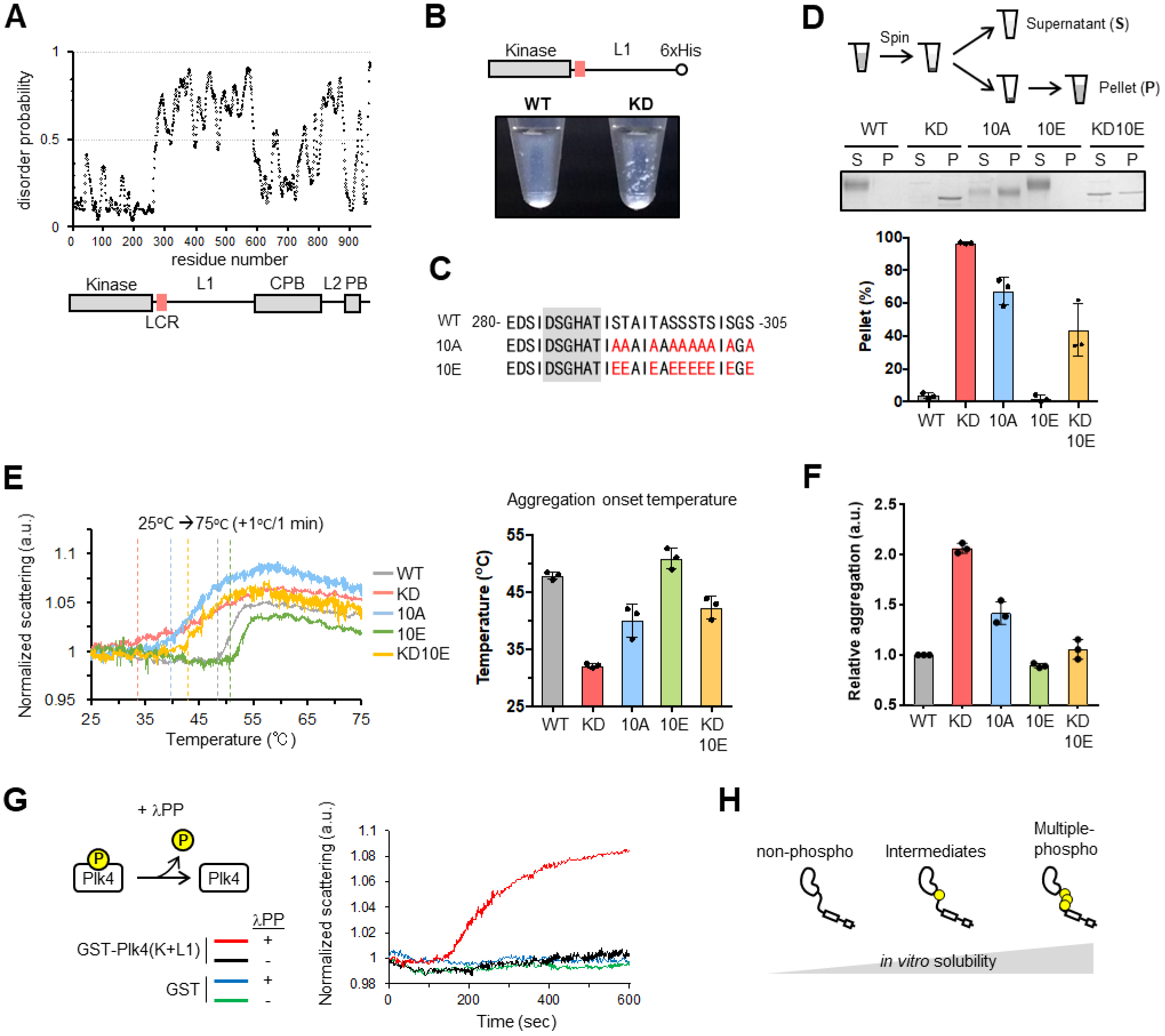
Plk4 modulates its solubility by auto-phosphorylation *in vitro*. **(A)** Disorder prediction and schematic of human Plk4 domains. Kinase, kinase domain; L1/L2, Linker 1/2; LCR, low-complexity region; CPB, Cryptic Polo-box; PB, Polo-box. **(B)** Image of 500 nM Plk4(Kinase+L1)-His6 fragments after GST-tag cleavage by PreScission protease. **(C)** Amino acid sequence of low complexity region (288-303 a.a.) and its neighboring region of human Plk4. Red letters, mutation sites; Gray background, degron motif. **(D)** Spin-down assay. After GST-tag cleavage, 500 nM Plk4(Kinase+L1)-His6 solution was centrifuged and separated into supernatant (S) and pellet (P) fractions. Representative gel stained with CBB and the quantification are shown. Graph represents mean±SD percentages of the pellet from three independent experiments. NaCl, 500 mM. **(E)** Measurement of the light scattering of GST-Plk4(Kinase+L1)-His6 (100 (μg/ml) under thermal control. Left: A representative data of three independent experiments. Values were normalized to the scattering at 25°C. Dotted vertical lines indicate each aggregation onset temperature. Right: Aggregation onset temperature. Graph shows mean±SD temperature of three independent experiments. **(F)** Measurement of protein aggregation of 300 nM GST-Plk4(Kinase+L1)-His6 by PROTEOSTAT aggregation assay. Fluorescence values were normalized to the WT. Graph shows mean ± SD of three independent experiments. **(G)** Dephosphorylation assay. GST-His6 or GST-Plk4(Kinase+L1)-His6 (100 (μg/ml) was incubated with λPP at 30 °C for the indicated time and the light scattering was measured. A representative data of three independent experiments is shown. **(H)** Schematic of biochemical property of Plk4.

To assess the solubility of purified Plk4 fragments *in vitro*, we performed a spin-down assay and showed that Plk4 WT fragments were mostly soluble, whereas the KD mutant was largely insoluble (Fig. 1D, S1C and S1D). As expected, the 10A mutant of the Plk4 fragment was mainly detected in the insoluble fraction, unlike the WT. In contrast, incorporation of the 10E mutation in the KD mutant (KD10E) improved the solubility of the Plk4 fragments compared with that of the KD mutant. We also monitored the light scattering of purified GST-Plk4(Kinase+L1)-His_6_ in solution as an indicator of protein aggregation under thermal control. When the temperature was increased, all the Plk4 fragments increased their light scatter but exhibited the onset of scattering at different temperatures (Fig. 1E and S1E). Compared with the WT, the KD and 10A mutants started scattering at lower temperatures. Adding the 10E mutation to the KD mutant shifted the onset of scattering to a higher temperature, than that of the KD mutant. In addition, dephosphorylation of the WT fragment by adding λ-phosphatase (λPP) significantly increased the scattering over time (Fig. 1G), demonstrating that phosphorylation of Plk4 prevents Plk4 aggregation. Furthermore, we confirmed the aggregation properties of Plk4 *in vitro* using the Proteostat dye, which fluoresces after being incorporated into protein aggregates. Similarly, KD, 10A and λPP-treated WT Plk4 fragments showed considerably greater protein aggregation compared with WT and the other mutants (Fig. 1F and S1F). Because it appeared that the 10A and 10E mutations did not completely mimic the KD and constitutively phosphorylated states of Plk4, respectively, it is possible that phosphorylation at other sites including the degron motif may also cooperatively regulate the solubility of Plk4 proteins. Taken together, we conclude that autophosphorylation of Plk4 regulates not only its protein degradation in cells but also its solubility by modulating its self-aggregating properties (Fig. 1H).

Recently, it has been shown that biomolecular condensates exhibit macromolecular structures such as spheres, networks and fibrils which are linked to their physiological functions (*25*, *26*). To characterize Plk4 self-assembly further, we visualized Plk4 fragments (Kinase+L1) and full-length Plk4 by labeling with GFP and mScarlet I, respectively. Intriguingly, we found that the KD mutant of GFP-Plk4 fragments (Kinase+L1) forms nano- to micrometer-scale, sphere-like aggregates *in vitro*, although such structures were barely detectable for the WT in this condition (Fig. 2A). This was also true for the full length Plk4 WT and KD (Fig. 2B). Interestingly, continuous incubation at 37 °C promoted self-assembly of the 10A mutant of the GFP-Plk4 fragment and further assembly of the KD mutant aggregates (Fig. 2A). This observation implies that Plk4 aggregates fuse to form larger aggregates, like protein condensates. These results demonstrate that in its nonphosphorylated state, Plk4 has an intrinsic ability to self-assemble to form condensate-like aggregates *in vitro*.

**Fig. 2.**
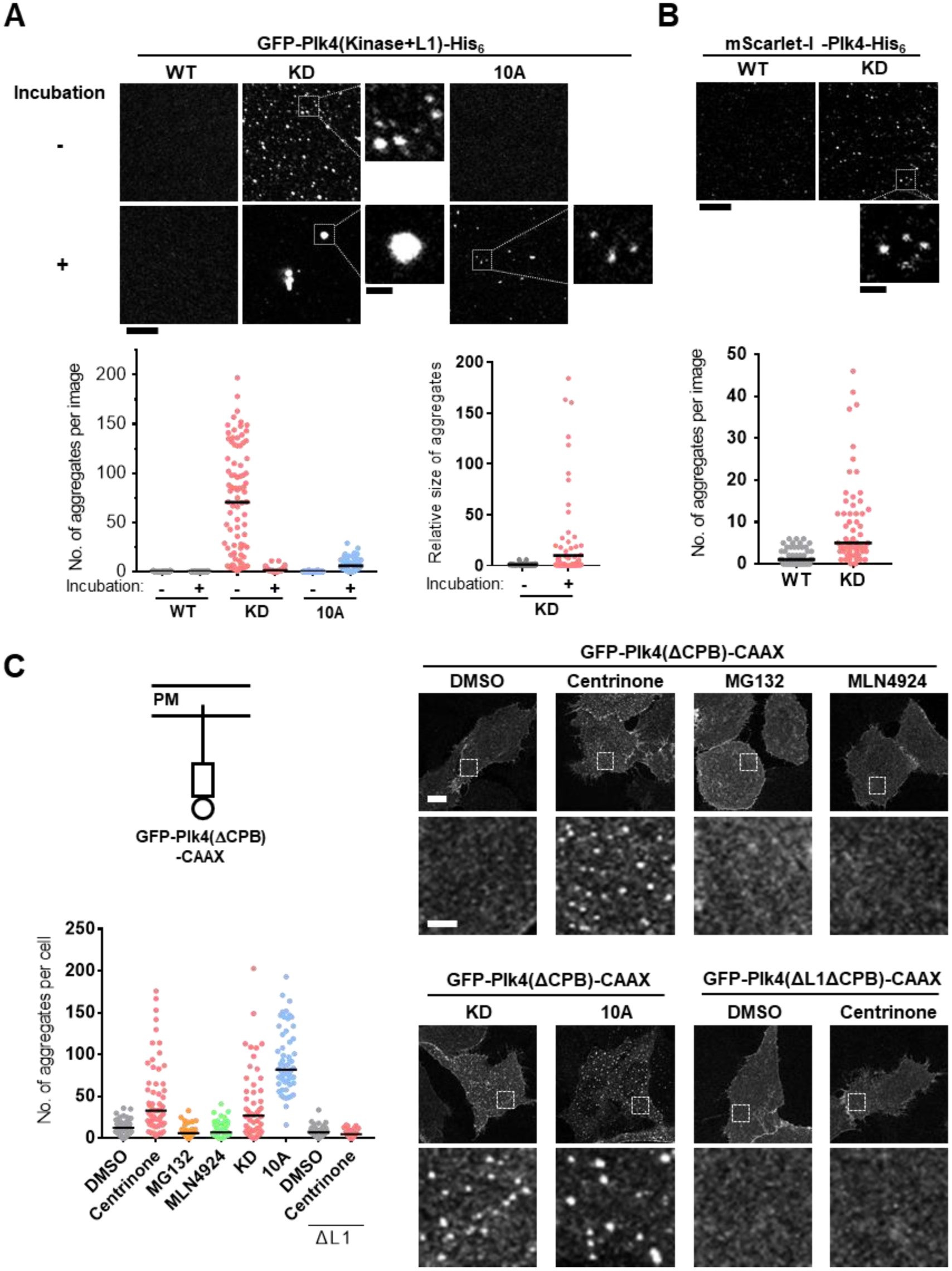
Plk4 forms condensate-like aggregates *in vitro* and in cells. **(A)** Fluorescence images of 100 nM GFP-Plk4(Kinase+L1)-His6 with or without incubation at 37 °C for 1 hour. Scale bar, 5 μm; inset, 1 μm. Left graph shows number of aggregates per image with median (black bar) of >60 images from three independent experiments. Right graph shows relative size of aggregates and the mean values (Black bar) of >140 aggregates. Data were normalized to the average size of “0 min”. **(B)** Fluorescence images of 50 nM mScarlet-Plk4-His6. Scale bar, 5 μm; inset, 1 μm. Graph shows number of aggregates per image with median (black bar) of >60 images from three independent experiments. **(C)** GFP-Plk4 (ΔCPB)-CAAX distribution at the plasma membrane region of HeLa cells stained with anti-GFP antibodies. Lower panels are magnified images of the indicated region of each cell. Cells were treated with DMSO, 100 μM centrinone (20 hours), 10 μM MG132 (5 hours) or 10μM MLN4924 (5 hours) and fixed 20 hours after plasmid transfection. Graph shows number of aggregates per plasma membrane region with median (black bar) of >50 cells from two independent experiments. Scale bar, 10 μm; 2 μm in magnified images.

We next asked whether Plk4 also self-assembles in cells. We next sought to evaluate solely the properties of Plk4 in cells and to exclude the effect of centrosome components, including those that interact with Plk4, on the behavior of Plk4. To this end, we designed a membrane-tethering system in which Plk4 lacking a CPB domain that binds to Cep152/Cep192 (*27*) and fused with a CAAX-motif localizes to the plasma membrane (Fig. 2C and S2A). When the Plk4 mutant was tethered to the plasma membrane in HeLa cells, known Plk4 interactors were undetectable at the site of the membrane-tethered Plk4 (Fig. S2B). Therefore, we considered that this system was suitable for specific visualization of Plk4 self-assembly in cells. Using this system, we found that GFP-Plk4(ΔCPB)-CAAX uniformly localized to the plasma membrane (Fig.2C and S3). In stark contrast, inhibition of the Plk4 kinase activity with centrinone, a specific inhibitor of Plk4, or overexpression of the KD and 10A mutants, significantly induced aggregation of membrane-localized Plk4 (Fig. 2C). Such aggregation was unlikely to be because of inhibition of autophosphorylation-dependent protein degradation, because inhibition of the degradation pathway by treatment with MG132 (an inhibitor of proteasomes) or MLN4924 (an inhibitor of the NEDD8-activating enzyme that activates SCF-cullin-RING ubiquitin ligase) did not influence the uniform localization of membrane-localized Plk4 (Fig. 2C). Furthermore, deletion of L1 (ΔL1) suppressed centrinone-induced aggregate formation, suggesting that a disordered L1 domain is required for the aggregation of Plk4 in cells (Fig. 2C). Overall, we conclude that Plk4 preferentially self-assembles in its nonphosphorylated state both *in vitro* and in cells.

### Regulated self-assembly mediates changes in centriolar dynamics of Plk4

What is the significance of Plk4 self-assembly for its localization and function at centrioles? To address this question, we first examined whether self-assembly regulates centriolar Plk4 dynamics by performing fluorescence recovery after photobleaching (FRAP) analysis with HeLa cells expressing GFP-Plk4. In control cells, overexpressed GFP-Plk4 showed dynamic turnover at centrioles (Fig. 3A). Intriguingly, we found that inhibition of Plk4 kinase activity with centrinone, but not inhibition of its protein degradation pathway by MG132 or MLN4924 treatment, significantly impaired the Plk4 turnover at centrioles (Fig. 3A), although all of these inhibitors similarly increased the amount of endogenous Plk4 at centrioles (Fig. 3B). These data suggest that centriolar Plk4 dynamics are regulated by its kinase activity. In other words, dissociation/diffusion of Plk4 from centrioles seems to be dependent on its kinase activity rather than on local protein degradation. Next, we tested whether introducing nonphosphorylatable mutations to Plk4 affected Plk4 turnover at centrioles (Fig. 3C). Consistent with the effects of the inhibitor treatment (Fig. 3A), KD, 10A and 13A mutations, but not a 2A mutation (in the degron motif), attenuated Plk4 turnover at centrioles (Fig. 3D and S3A). Together with the *in vitro* data, these results indicate that the dynamics of Plk4 at centrioles changes depending on autophosphorylation-regulated self-assembly. We speculate that the autophosphorylation-regulated self-assembly properties of Plk4 may be conserved across species, because *Drosophila* Plk4 (Dm Plk4) exhibited similar behavior in HeLa cells (Fig. S3C-F).

**Fig. 3.**
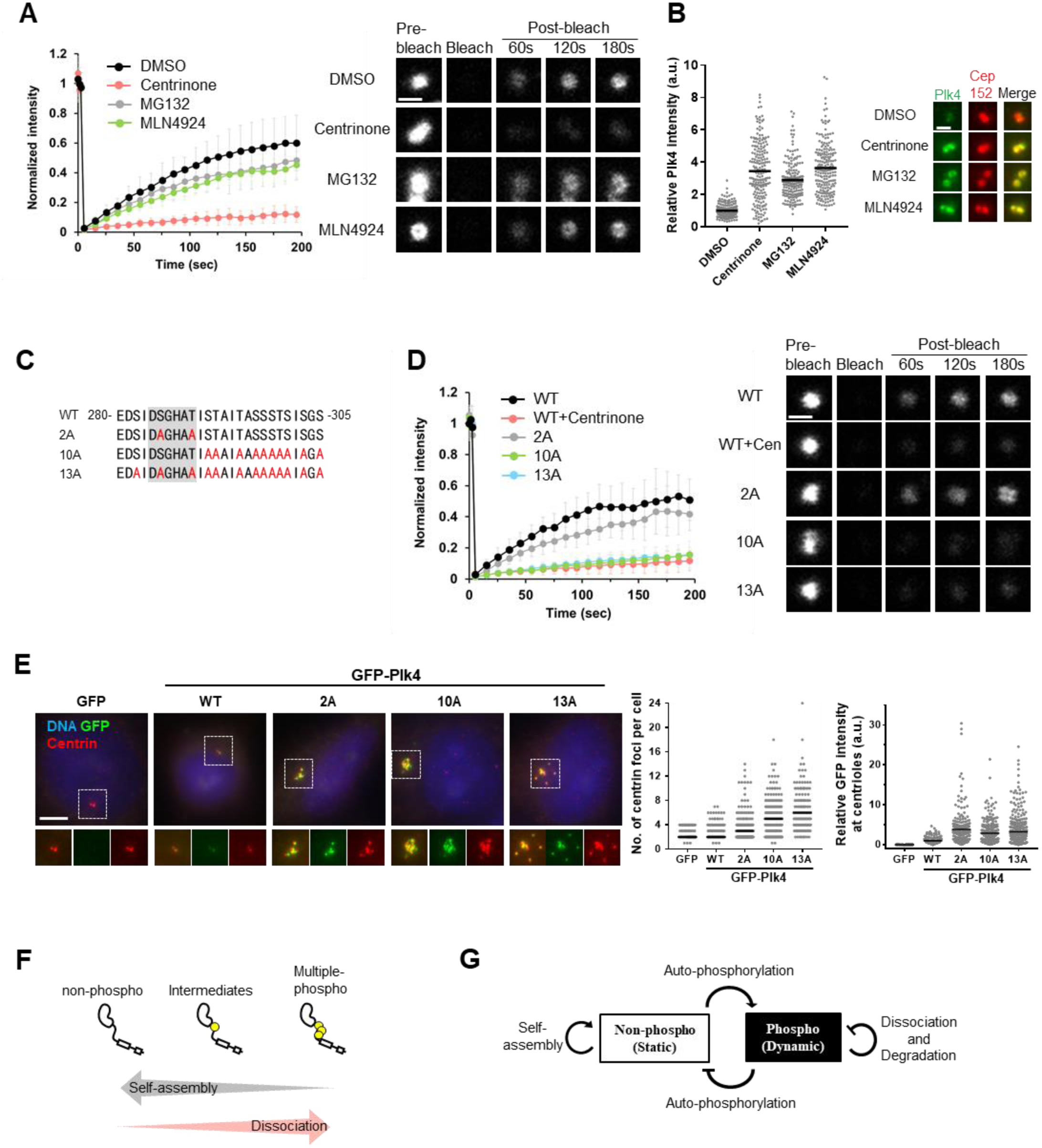
Autophosphorylaiton controls centriolar Plk4 dynamics by regulating self-assembly. **(A)** FRAP analysis of GFP-Plk4 in HeLa cells. Cells were treated with DMSO, 100 μM centrinone, 10 μM MG132 or 10 μM MLN4924 for 5-6 hours respectively. For **(A)** and **(D)**, intensities were normalized with the average of three pre-bleach signals. Graph shows mean±SD of 15 cells from three independent experiments. **(B)** Effects of inhibitors on endogenous centriolar Plk4 in HeLa cells. Cells were stained with the indicated antibodies and centriolar Plk4 intensity was quantified. Graph shows signal intensity of endogenous Plk4 at centrioles. Black bars show mean values of >90 cells from three independent experiments. Data were normalized to the average intensity of DMSO. **(C)** Amino acid sequence of mutation sites in Plk4. Red letters, mutation sites; Gray back ground, degron motif. **(D)** FRAP analysis of GFP-Plk4 mutants in HeLa cells. **(E)** Centriole over-duplication assay. GFP-Plk4 was exogenously expressed in HeLa cells under human Plk4 promoter. Cells were fixed 48 hours after plasmid transfection and stained with the indicated antibodies. Scale bar, 5 μm; inset, 1 μm. Left graph represents the number of Centrin foci in GFP-positive interphase cells. Black bar shows median of >100 cells from three independent experiments. Left graph represents GFP signal intensity at centrioles. Data were normalized to the average intensity of “WT”. Black bar shows mean signal intensity of >80 cells from two independent experiments. **(F)** Schematic summary of Plk4 property *in vitro* and at centrioles. **(G)** Proposed self-organization system of Plk4.

We next asked whether the regulated self-assembly of Plk4 is involved in proper centriole duplication. In human cells, exogenous overexpression of Plk4 under a strong promoter such as CMV (cytomegalovirus) induces centriole overduplication (*5*, *16*). To express GFP-Plk4 mutants in HeLa cells at levels comparable to endogenous expression, we utilized the endogenous promoter of human Plk4 (*28*). In this condition, expression of GFP-Plk4 WT resulted in minimal centriole overduplication (Fig. 3E and S3B). However, the 2A mutation in the degron motif of Plk4 led to significant centriole overduplication, presumably by increasing Plk4 expression. Importantly, 10A and 13A mutations induced over-duplication of centrioles, to greater extent than the WT and the 2A mutant (Fig. 3E and S3B). Although the 10A and 13A mutations may also affect the protein degradation of Plk4 to some extent, these results suggest that to ensure the proper centriole copy number, Plk4 self-assembly must be controlled by autophosphorylation. Overall, we conclude that autophosphorylation of Plk4 promotes not only its degradation but also its dissociation/diffusion dynamics at centrioles by regulating self-assembly and also that this property of Plk4 is critical for proper control of the centriole copy number (Fig. 3F).

Based on previous knowledge and our current results, we propose a self-organization model comprising two components, phosphorylated and nonphosphorylated-Plk4 species (Fig. 3G). This study described the mutual interactions of these two components, which are reminiscent of the reaction-diffusion system represented by the Turing model that can be applied to a variety of spatial pattern formation (*29*, *30*). Similarly, the self-organization model may explain how Plk4 can form spatial patterns at centrioles by means of its dynamic properties including self-assembly and dissociation/diffusion.

### Autophosphorylated Plk4 appears as a single focus at mother centrioles prior to STIL-HsSAS6 loading

Our results thus far raised the possibility that phosphorylated and nonphosphorylated Plk4 may cooperatively regulate their distribution patterns around mother centrioles by self-organization. Consistent with this concept, we reported that Plk4 forms a biased ring in G1 phase before centriolar loading of STIL-HsSAS6 (*9*, *31*). We first confirmed that in G1 phase, Plk4 formed a ring-like structure with an intense focus (Fig. 4A and B). Thereafter, Plk4 changed its localization pattern to a single focus that colocalized with STIL and HsSAS6 in G1/S phase, as reported previously (*9*, *32*) (Fig. 4A and B). Next, to monitor autophosphorylation of Plk4 in cells, we newly generated the antibody against phosphorylated serine 305 (pS305) of Plk4 in this study (*19*, *31*) (Fig. S4A-D). Strikingly, structural illumination microscopy (SIM) revealed that even in cells with a ring-like distribution of Plk4, Plk4pS305 appeared as a single focus around mother centrioles (Fig. 4B). In addition, the Plk4pS305 signal overlapped with the intense focus of Plk4 rings (Fig. 4B and S5A). Consistent with this, even in cells in which HsSAS6 had not yet been loaded to the centrioles, the Plk4pS305 signals were already patterned as a single focus around mother centrioles (Fig. 4C). After HsSAS6 loading, the Plk4pS305 signals colocalized with HsSAS6 as a focus (Fig. 4C). Thus, these results demonstrate that, before procentriole formation, distribution of autophosphorylated Plk4 is already biased around mother centrioles, which could provide the assembly site of procentrioles. As reported previously, centriolar Plk4pS305 signals increased after HsSAS6 loading, suggesting further activation of Plk4 via STIL-HsSAS6 loading (Fig. 4D and S4E). In addition, even in cells depleted of HsSAS6 (siHsSAS6-treated), the Plk4pS305 signal was localized as a single focus around mother centrioles in S-phase (Fig. 4E), further confirming that the occurrence of Plk4 phosphorylation is independent of the presence of HsSAS6 and is likely dependent on the intrinsic properties of Plk4.

**Fig. 4.**
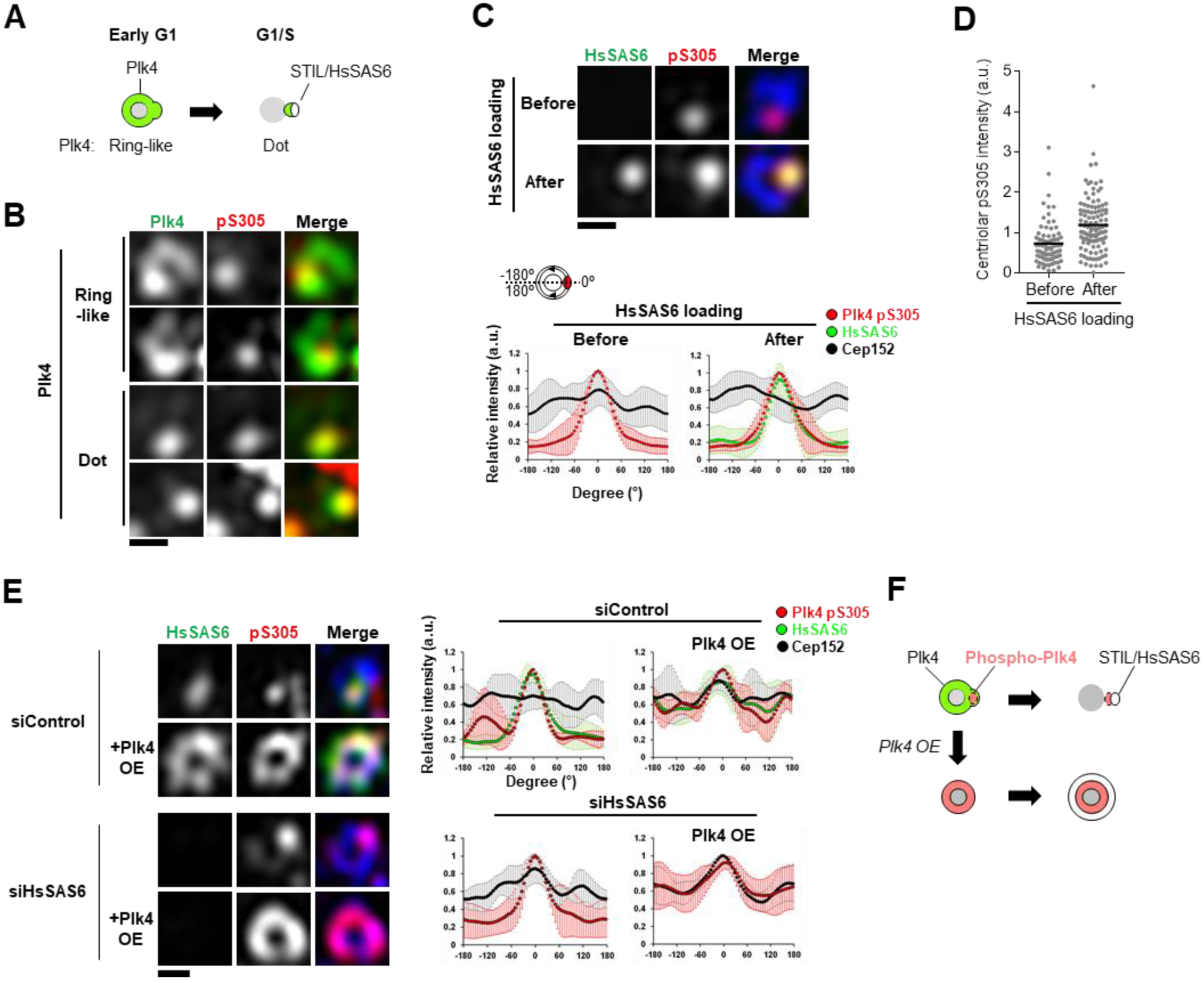
A single focus of auto-phosphorylated Plk4 is generated prior to STIL-HsSAS6 loading. **(A)** Schematic of Plk4 distribution around mother centriole in centriole duplication. **(B)** 3D-SIM images of centrioles immunostained with the indicated antibodies in HeLa cells. Cep152 was costained and used for a marker of mother centriole wall. In merged images, green and red represent Plk4 and Plk4pS305, respectively. Scale bar, 0.3 μm **(C and E)** 3D-SIM images of centrioles immunostained with antibodies against HsSAS6, Plk4pS305 and Cep152 in HeLa cells. In merged images, green, red and blue represent HsSAS6, Plk4pS305 and Cep152, respectively. Relative intensities of each protein were quantified along Cep152 ring and maximum peaks of Plk4pS305 intensity were set to 0°. Intensities were normalized to each maximum intensity. Graph shows mean ±SD. Scale bar, 0.3 μm **(C)** Plk4pS305 patterns before and after HsSAS6 loading. Each n = 10 cells from two independent experiments. **(D)** Quantification of Plk4pS305 intensity at centrioles before and after HsSAS6 loading. HsSAS6 loading “Before or After” was distinguished by setting a threshold of centriolar HsSAS6 intensity (See Figure S4E). Graph shows mean values (Black bars) of centriolar Plk4pS305 intensities of 100 cells from two independent experiments. **(E)** Plk4pS305 patterns in S-phase arrested HeLa cells. Cells were treated with siRNA for 48 hours and 6 μM aphidicolin for 24 hours. For Plk4 overexpression (OE), cells were transfected with plasmids encoding Plk4-3×FLAG and fixed 24 hours after plasmid transfection. n = 5 cells (for siControl) or n = 10 cells (for siHsSAS6) from two independent experiments. **(F)** Schematic of phospho-Plk4 (pS305) patterns.

To examine whether this spatial pattern of Plk4pS305 is generated through self-organization, we perturbed the putative self-organization system by overexpressing Plk4. Overexpression of Plk4 interrupted the formation of a single focus of Plk4pS305 and instead induced formation of a uniform ring of Plk4pS305 around mother centrioles in both siControl and siHsSAS6-treated cells (Fig. 4E). Thus, these results suggest that endogenous expression levels of Plk4 self-organize autophosphorylated Plk4 to a single focus around mother centrioles, which presumably limits STIL-HsSAS6 loading to a single focus (Fig. 4F).

### Autonomous activation of Plk4 drives centriolar loading of STIL-HsSAS6

Our results demonstrated that Plk4 is autophosphorylated before STIL-HsSAS6 loading, suggesting the possibility that autonomous activation of Plk4 is a consequence of Plk4 self-assembly and drives centriolar loading of STIL-HsSAS6 for the initiation of procentriole formation. To address these possibilities, we monitored the time course of Plk4 autophosphorylation and STIL-HsSAS6 loading in cells transiently treated with centrinone at G1 phase. We first inhibited the kinase activity of Plk4 by centrinone treatment in G1-arrested HeLa cells and subsequently released the inhibition by washout of centrinone (Fig. 5A). We found that transient centrinone treatment was sufficient to remove the Plk4pS305 signal from centrioles, whereas Plk4 itself significantly accumulated at centrioles (Fig. 5B-E). As previously reported (*16*), centrinone treatment induced delocalization of STIL-HsSAS6 from centrioles, indicating that Plk4 kinase activity is required for centriolar localization of the STIL-HsSAS6 complex (Fig. 5B-E). After washout of centrinone, accumulation of Plk4pS305 and STIL-HsSAS6 signals at centrioles began almost simultaneously and increased gradually in most cells (Fig. 5B-F). Importantly, even in HsSAS6-depleted cells (siHsSAS6-treated), the autophosphorylated Plk4 (pS305) signal was increased after washout of centrinone, as observed in control cells (siControl-treated) (Fig. 5B, D and S5C), indicating that autophosphorylation of Plk4 occurs in Plk4 aggregates independently of centriolar loading of the STIL-HsSAS6 complex. In contrast, as has been reported previously, depletion of HsSAS6 suppressed centriolar loading of STIL (Fig. 5C and E). Taken together, our results suggest that (1) autonomous activation of Plk4 within the Plk4 ring occurs in G1 phase, (2) Plk4 activation is mediated by its self-assembly properties, and (3) Plk4 activation subsequently drives centriolar loading of the STIL-HsSAS6 complex by phosphorylating STIL.

**Fig. 5.**
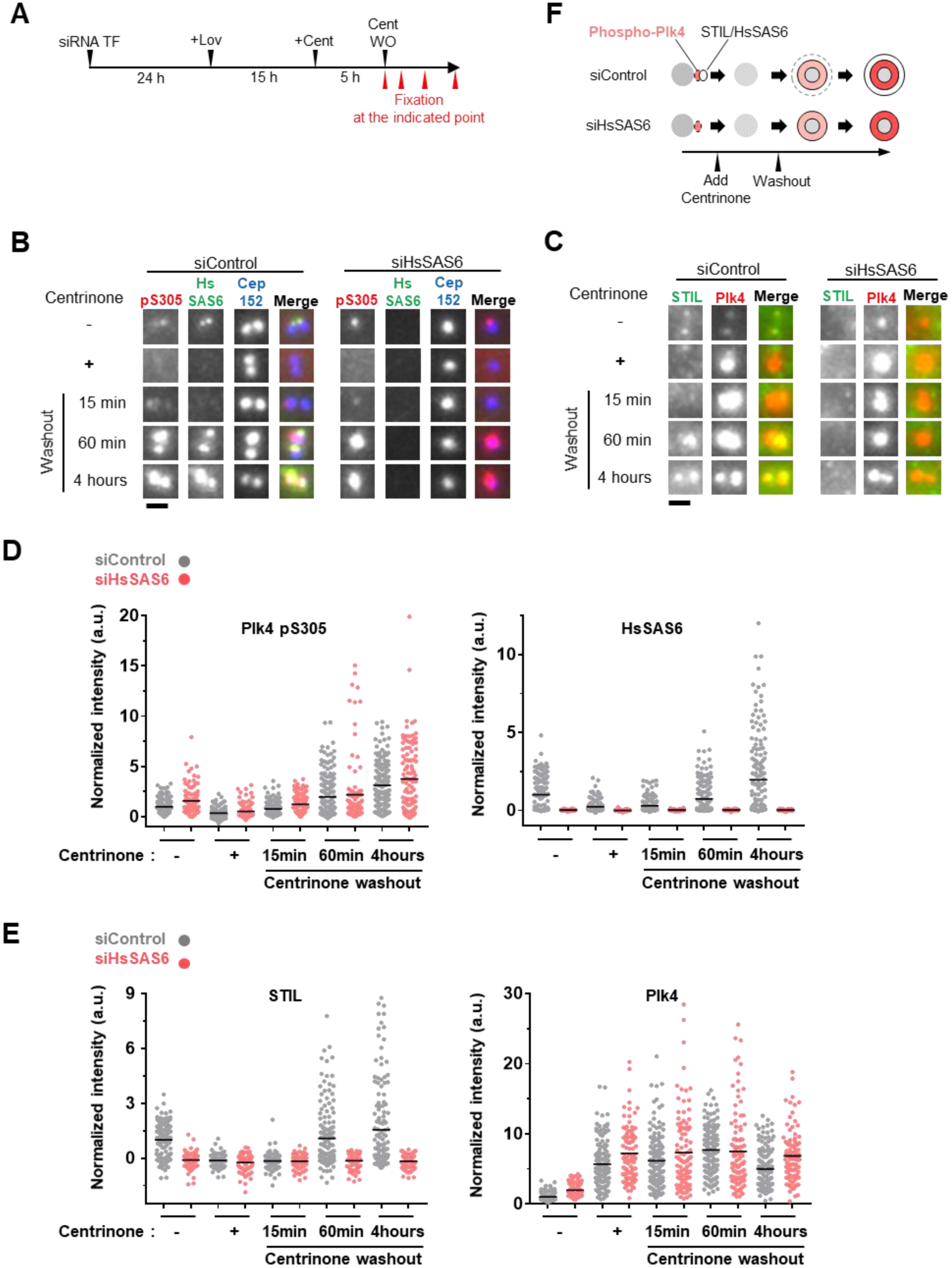
Autonomous activation of Plk4 that stems from Plk4 self-assembly drives centriolar loading of STIL-HsSAS6. **(A)** Scheme of the experimental condition. G1-arrested HeLa cells (10 μM Lovastatin (Lov), 20 hours (See Figure S5B)) were treated with 100 μM centrinone (Cent) and then washed out (WO) the centrinone. **(B)** Representative images of immunostained HsSAS6 and Plk4pS305 in the indicated condition. Scale bar, 1 μm. **(C)** Representative images of immunostained STIL and Plk4 in the indicated condition. Scale bar, 1 μm. **(D and E)** Quantification of centriolar Plk4pS305, HsSAS6, STIL and Plk4 intensities in the indicated condition. Gray and pink dots represent the data from siControl and siHsSAS6-treated cells, respectively. Black bars show mean values of >80 cells from two independent experiments. Intensities were normalized to the average intensity of “siControl & Centrinone(-)” condition. **(F)** Schematic summary of the correlation between centriolar phospho-Plk4 (pS305) and STIL-HsSAS6 intensity in Figure 5 B-E.

## Discussion

In this study, we demonstrated that autophosphorylation of Plk4 modulates its self-assembly and dissociation/diffusion dynamics both *in vitro* and in cells. From these findings, we hypothesize that the intrinsic properties of Plk4 are responsible for its spatial distribution. In our model (Fig. 6), phosphorylated and nonphosphorylated Plk4 species that have different diffusion coefficients interact with each other to generate the biased pattern of Plk4. The interaction mode of these two components is similar to Turing’s reaction-diffusion model. In fact, autophosphorylated Plk4 is spatially distributed as a single focus around mother centrioles. We thus propose that Plk4 makes a kinase-active focus around the mother centriole through self-organization, which provides the single site for the recruitment of STIL-HsSAS6. Furthermore, given that Plk4 is activated by trans-autophosphorylation depending on its local concentration (*14*), we assume that increasing amounts of centriolar Plk4 from late mitosis to late G1 phase could be a trigger for autonomous activation (*32*, *34*). In our model, the most important issue to address is how the Plk4 self-organization system generates only one focus of active Plk4 around the mother centriole. To investigate and improve our model further, other approaches such as super-resolution live imaging and mathematical simulation analyses should be applied.

**Fig. 6.**
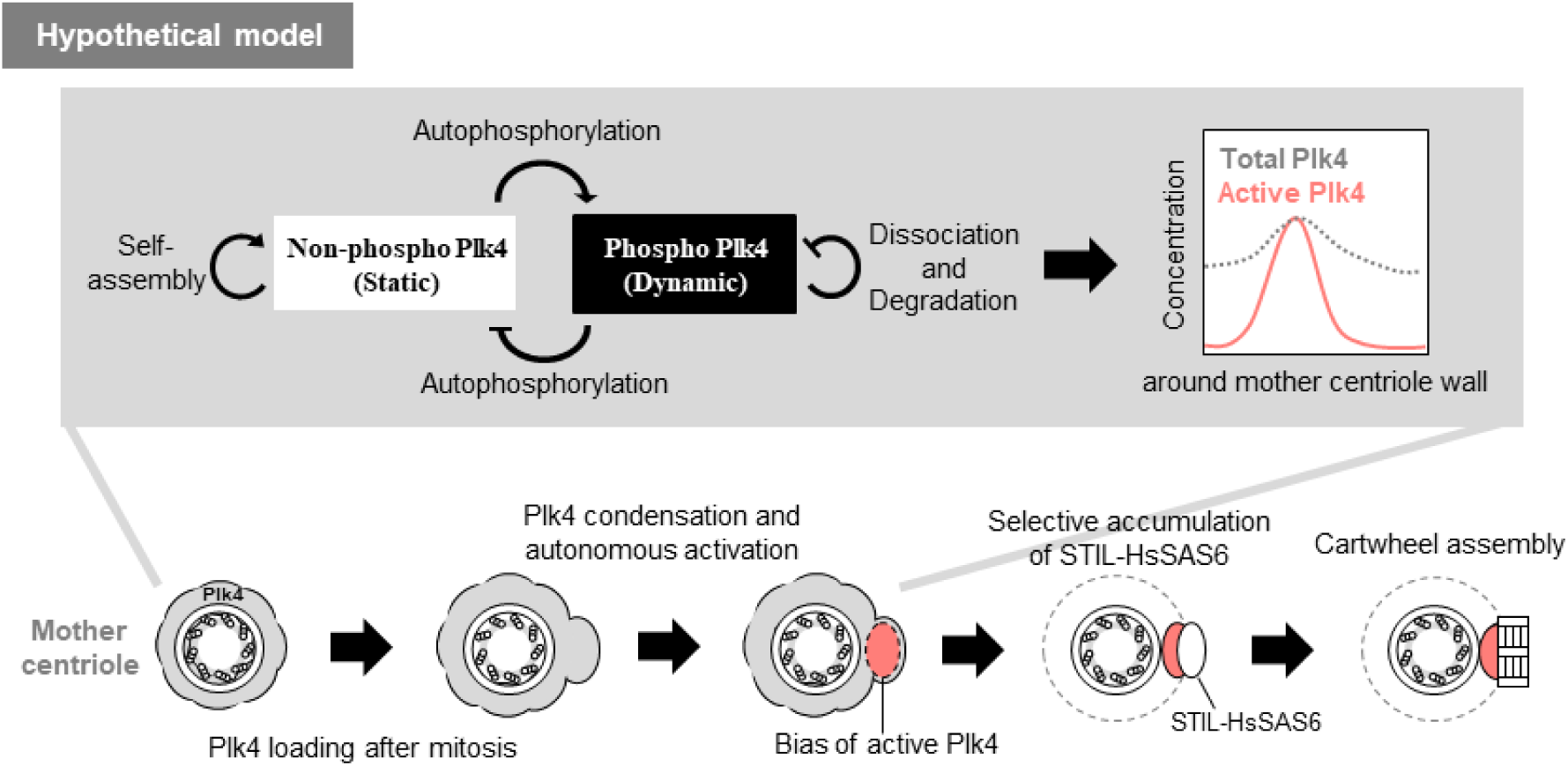
A hypothetical model: Self-organization of Plk4 directs a single daughter centriole formation per a mother centriole. Schematic illustration of a hypothetical model. Increasing centriolar Plk4 level after mitosis drives condensation and autonomous activation of Plk4 prior to STIL-HsSAS6 loading. Self-organization property of Plk4 induces spatial pattern formation of autophosphorylated Plk4, which results in a single focus formation of activated Plk4. The single focus is a possible target site for STIL-HsSAS6 loading.

Autophosphorylation of Plk4 triggers its own degradation via the SCF ubiquitin ligase-proteasome pathway (*15*, *16*, *18*, *20*). Indeed, we observed that inhibition of SCF ubiquitin ligase or proteasome increased the amount of Plk4 at centrioles. However, our FRAP analysis suggest that centriolar Plk4 dynamics are predominantly regulated by self-assembly via autophosphorylation rather than by protein degradation; i.e., by dissociation of Plk4 from centrioles. Because it is debatable whether ubiquitin-proteasome machineries can directly ubiquitinate and degrade centriolar Plk4 in a PCM-crowded centrosome, we speculate that the degradation machinery may mainly control the cytoplasmic pool of Plk4, whereas the amount of centriolar Plk4 is controlled by autophosphorylation-regulated self-assembly and dissociation. In addition, we reason that autophosphorylation-dependent dissociation/diffusion of Plk4 from centrioles could increase fidelity by restricting centriole duplication via more rapid delocalization of excess Plk4 from centrioles. In our experiments, the 10A mutation of Plk4 induced centriole overduplication (Fig. 3E and S3B). Given that previous studies have suggested that multiple phosphorylation within neighboring regions of the DSG degron somehow enhance Plk4 degradation (*16*, *20*), it is unclear whether centriole overduplication as a result of the 10A mutation of Plk4 results from defects in degradation of Plk4 or in defects of regulated self-assembly. Because it has been suggested recently that self-aggregation protects the protein from degradation (*23*), it is possible that degradation of Plk4 is also suppressed by its own aggregation in its nonphosphorylated state. Thus, future investigations are required to distinguish between the effects of autophosphorylation on Plk4 degradation and dissociation/diffusion.

Recent studies have revealed the intrinsically disordered region (IDR) and LCR act in protein condensation (*35*, *36*). Moreover, it has been shown that protein condensation can be regulated by post-translational modifications (*22*–*24*, *37*, *38*). We demonstrated that an IDR of Plk4 is required for its self-assembly. The Plk4 self-assembly is regulated by autophosphorylation within the LCR. Thus, our results suggest that Plk4 has a common feature on the regulation of protein condensation. Because protein condensates anchor and concentrate the interactors into the structure to allow its function (*26*, *39*), self-assembled Plk4 may function for selective accumulation of STIL-HsSAS6 to trigger formation of procentrioles.

Although our study mainly focused on Plk4 property before centriolar loading of STIL-HsSAS6, it remains to be elucidated how the focus of active Plk4 is retained as a focus around mother centrioles after STIL-HsSAS6 loading. We speculate that although Plk4 self-assembly seems to be released by autophosphorylation, stable interaction with STIL-HsSAS6 retains active Plk4 to the focus at mother centrioles. It will be important to clarify how such transition of Plk4 state is regulated by forming a complex with STIL-HsSAS6 at the procentriole assembly site. Furthermore, another important issue to address in the future would be the function of Cep152 and Cep192, known scaffolds for Plk4, in the regulation of Plk4 status. In particular, it would be conceivable that Cep152 regulates the activity and phosphorylation state of Plk4 on the mother centriole wall and modulates spatial pattern of Plk4 there. Further investigation of the biochemical nature of procentriole components will expand our molecular understanding of centriole duplication.

## Acknowledgements

We thank Yuka Nozaki and Tomoko Ashikawa for supporting experiments; Noriko Tokai for technical advice of 3D-SIM analysis; all the members of Kitagawa laboratory for discussions and critical reading of the manuscript. This work was supported by Grant-in-Aid for Young Scientists (A) and for JSPS Fellows and for Scientific Research on Innovative Area from the Ministry of Education, Science, Sports and Culture of Japan, by Takeda Science Foundation, by Mochida Memorial foundation.

## Author contributions

S.Y. and D.K. designed the study and the experiments. S.Y. performed all of the experiments. S.Y. and D.K. wrote the manuscript.

---

**Fig. S1.**
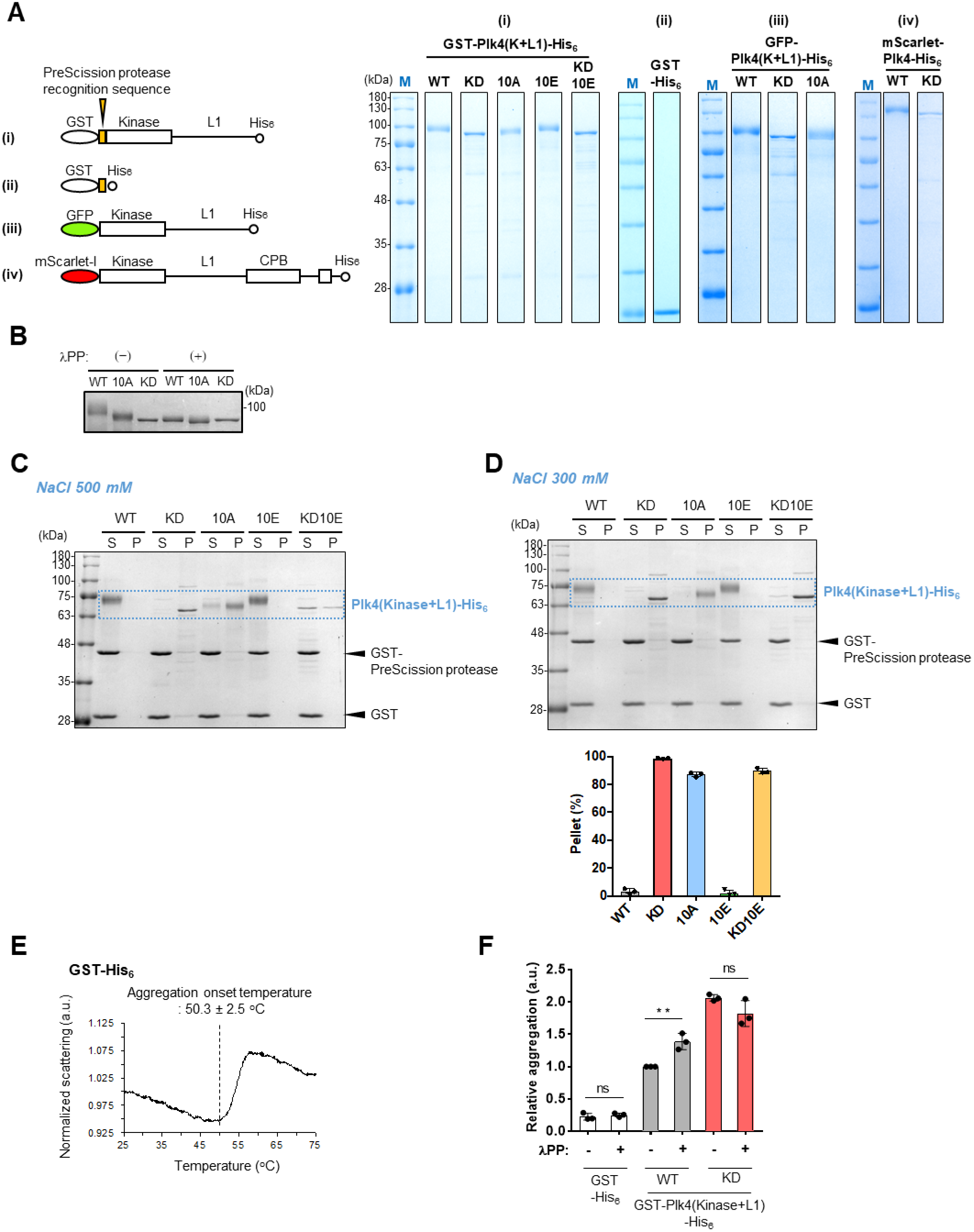
*in vitro* analysis of purified Plk4. **(A)** Purified proteins used in this study. Images show SDS-PAGE gel after CBB staining. “M”(Blue letter) indicates markers of molecular weight. **(B)** GST-Plk4 (Kinase+L1) fragments were dephosphorylated with λPP. Image shows CBB-stained gel. Band shift of WT was suppressed completely by KD mutation and λPP treatment, and largely by 10A mutation. **(C)** Original image of spin-down assay in Figure 1D (NaCl concentration, 500 mM). For (B) and (C), a dotted blue rectangle shows the bands of Plk4(Kinase+L1)-His_6_. Black arrowheads indicate the positions of GST-tagged PreScission protease and cleaved GST-tag. Importantly, GST-PreScission protease and GST-tag were mostly detected in the soluble fraction. **(D)** Spin-down assay in lower NaCl condition (300 mM). As in Figure 1D and S1C, 500 nM GST-Plk4 (Kinase+L1)-His_6_ solutions were centrifuged and separated into supernatant (S) and pellet (P) fractions, after GST cleavage. The graph shows mean percentages of pellet and SD from three independent experiments. **(E)** Measurement of light scattering of GST-His_6_ (100 (μg/ml). The graph is a representative data of three experiments. Data was normalized to the scattering at 25°C. Dotted vertical line indicates aggregation onset temperature. **(F)** PROTEO STAT aggregation assay with or without λPP. Fluorescence intensities were normalized to the “WT, λPP(-)”. The graph shows mean values and SD of three independent experiments. This experiment was performed at the same time of Figure 1F and the data of λPP(-) are the same as Figure 1F. **p<0.01, ns, not significant (Two-tailed t-test).

**Fig. S2.**
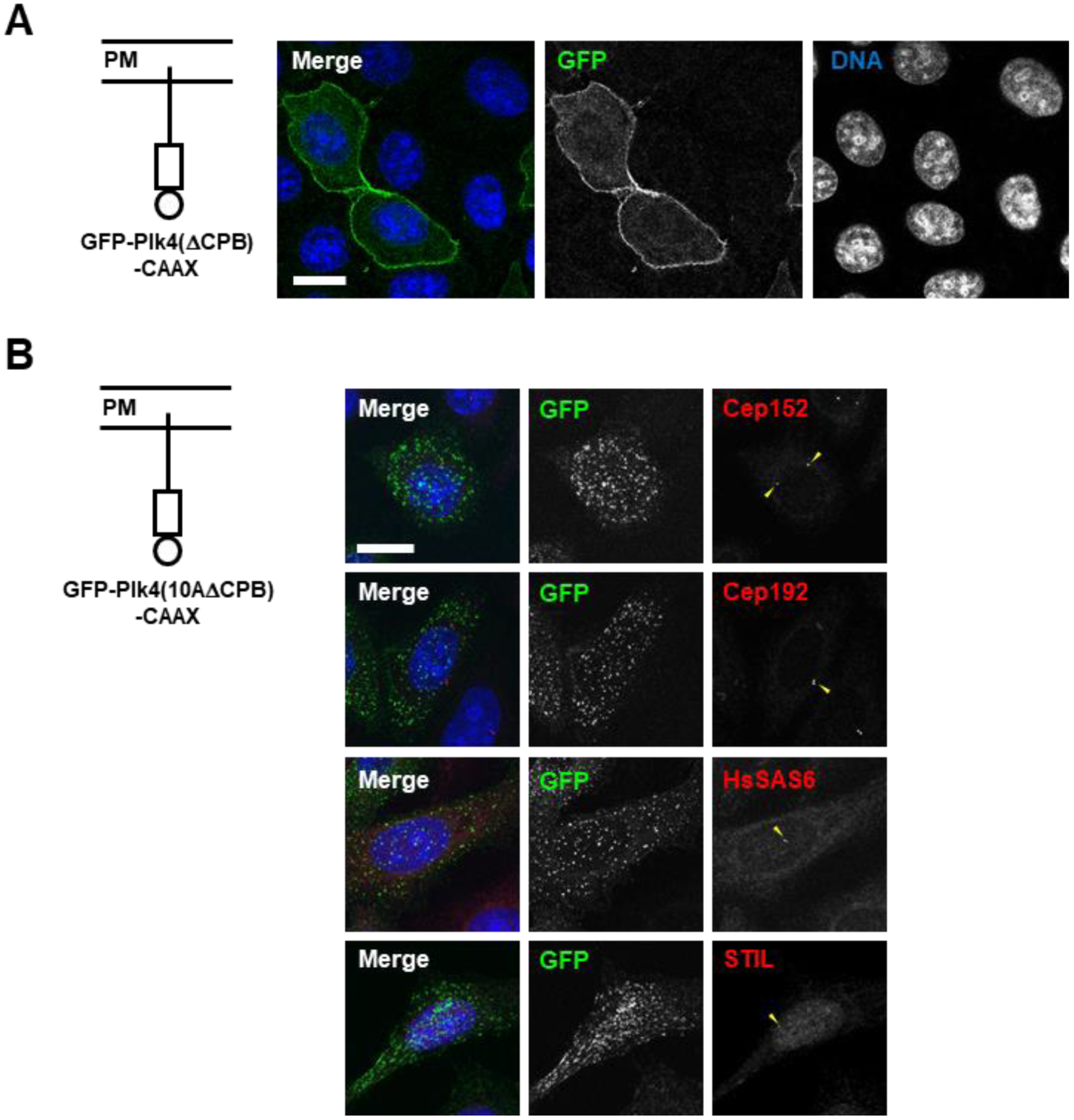
Characterization of GFP-Plk4-CAAX in HeLa cells. **(A)** Image of HeLa cells expressing GFP-Plk4 (ΔCPB)-CAAX. GFP-Plk4 (ΔCPB)-CAAX could be observed at the plasma membrane region. Image was obtained by immunostaining with anti-GFP antibodies. Scale bar, 15 μm. **(B)** Immunostaining of Cep152, Cep192, HsSAS6 and STIL in HeLa cells expressing GFP-Plk4(10AΔCPB)-CAAX. In this condition, each known interactor was undetectable at the foci of membrane-tethered Plk4. Yellow arrowheads indicate centrosomal signal of the indicated proteins. Scale bar, 15 μm.

**Fig. S3.**
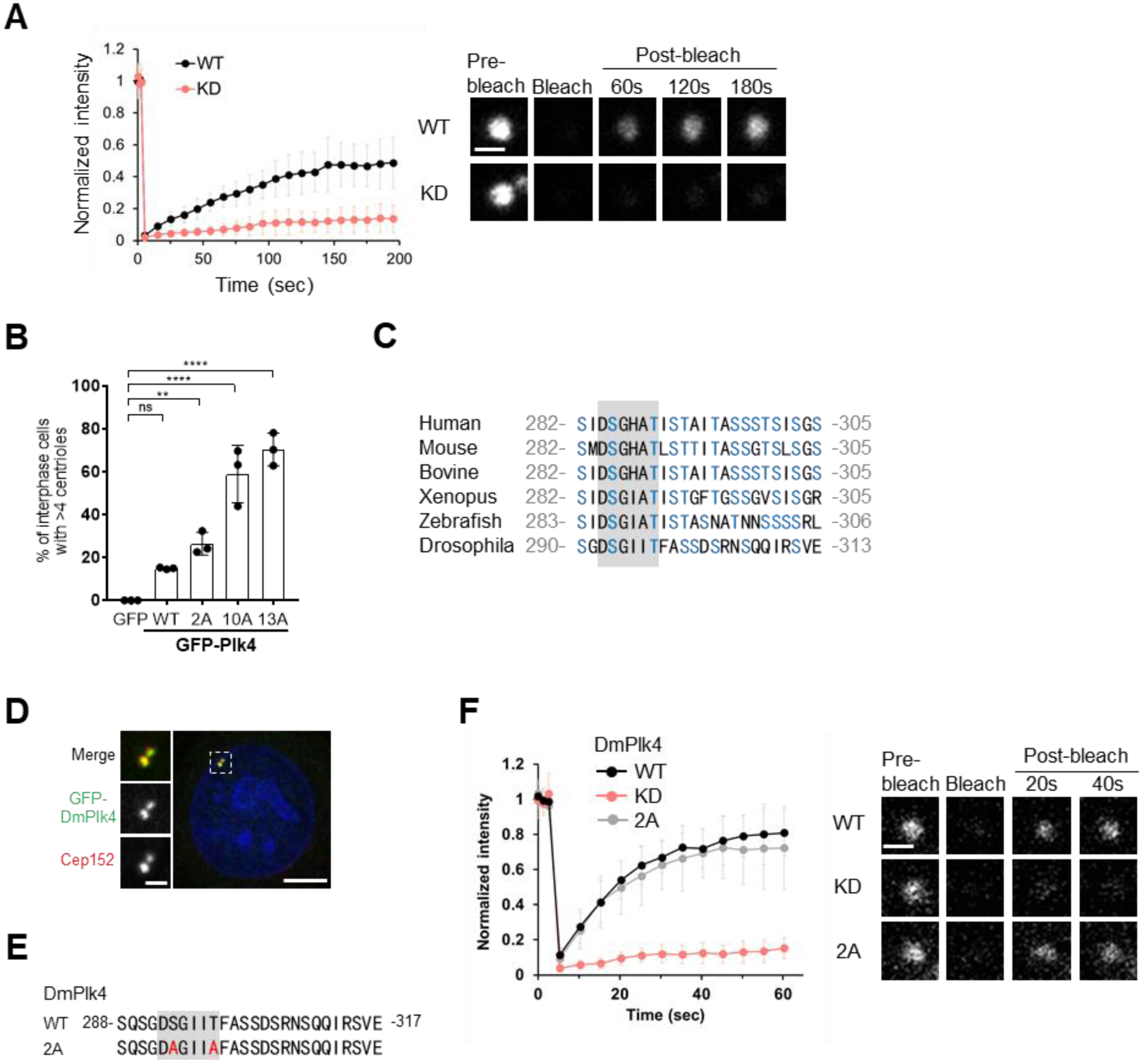
Analyses of Plk4 self-assembly at centrioles. **(A)** FRAP analysis of GFP-human Plk4 WT and KD overexpressing in HeLa cells. Intensities were normalized with the average of three pre-bleach signals. The graph shows mean values and SD of 15 cells from three independent experiments. Scale bar, 1 μm. **(B)** Centriole overduplication assay, related to Figure 3E. Percentages of interphase cells which have >4 centrin foci were calculated in the same experiments with Figure 3E. The graph represents mean percentages and SD of three independent experiments. \*\**p* < 0.01, \*\*\*\**p* < 0.0001, ns, not significant, one-way ANOVA. **(C)** Conservation of S/T enriched sequence of Plk4. Amino acid sequence alignment of Plk4s is shown. Gray background indicates degron. Serine and threonine are shown with blue letters. S/T enriched property around degron motif seems to be conserved across species. **(D)** HeLa cells expressing GFP tagged Drosophila Plk4 (DmPlk4). After transfection of the plasmid encoding GFP-DmPlk4, cells were fixed and stained with anti-GFP and Cep152 antibodies. Scale bar, 5 μm in larger image and 1 μm in magnified image. **(E)** Amino acid sequence of degron motif and the neighboring region in DmPlk4. Gray background indicates degron. **(F)** FRAP analysis of GFP-DmPlk4 overexpressing in HeLa cells. Intensities were normalized with the average of three pre-bleach signals. The graph shows mean values and SD of 15 cells from three independent experiments. Scale bar, 1 μm.

**Fig. S4.**
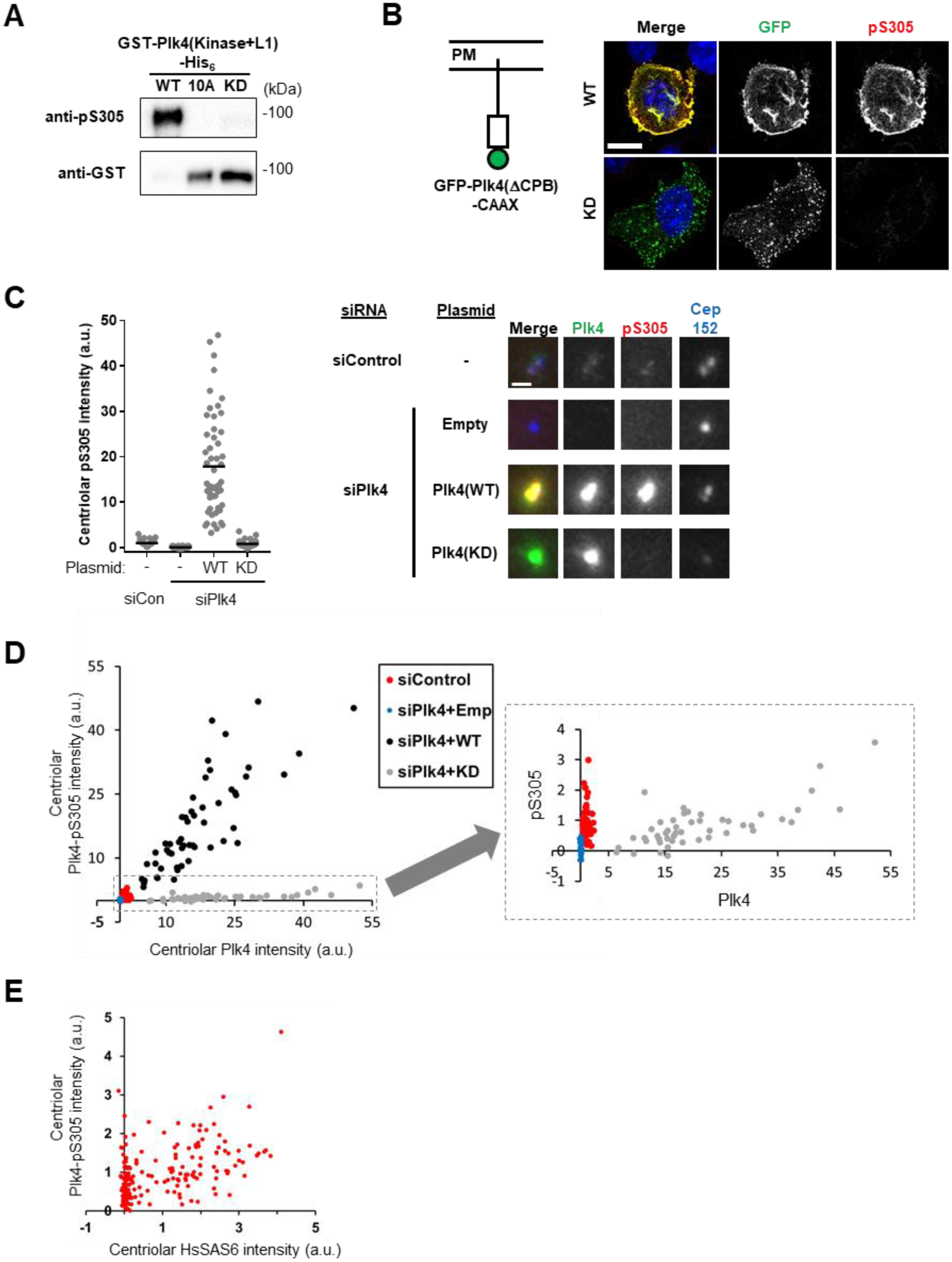
Validation of the specificity of anti-Plk4pS305 antibodies. **(A)** Immunoblotting of GST-Plk4(kinase+L1)-His_6_ with anti-Plk4pS305 (upper) and anti-GST (lower) antibodies. Anti-pS305 antibodies specifically reacted with purified Plk4 WT fragments, but hardly with 10A and KD. Because the band of WT was smeared probably by phosphorylation, detection of WT with anti-GST antibodies became weaker, compared with KD. **(B)** Immunostaining of Plk4pS305 in HeLa cells expressing GFP-Plk4(ΔCPB)-CAAX. Cells were fixed and stained with the indicated antibodies. Anti-pS305 antibodies specifically reacted with membrane-tethered Plk4 (WT), but hardly with KD. Scale bar, 15 μm. **(C)** Immunostaining of Plk4pS305 in HeLa cells transfected with the indicated siRNA and plasmids. Cells were fixed 48 hours after siRNA transfection and 24 hours after plasmid transfection. Plk4-3×FLAG vectors were used for Plk4 overexpression and 3×FLAG vectors were used for “empty” vector transfection. Left panel shows centriolar Plk4pS305 intensity. Data were normalized to the average intensity of siControl. The graph shows mean values (Black bar) of >45 cells from two independent experiments. Scale bar, 1 μm. **(D)** Correlation of centriolar Plk4 intensity and Plk4pS305 intensity in the same experiments with Figure S4C. Anti-pS305 antibodies reacted with centriolar Plk4. In the case of Plk4 overexpression, the antibodies showed high specificity against WT, compared with KD. Magnified graph (Right panel) represents the same data of "siControl", "siPlk4+Empty" and "siPlk4+Plk4WT". **(E)** Correlation of centriolar HsSAS6 and Plk4pS305 intensity in HeLa cells used in Figure 4D. Cells were costained with anti-HsSAS6 and anti-Plk4pS305 antibodies. The graph shows centriolar HsSAS6 and Plk4pS305 intensities of 100 cells from two independent experiments. Data were normalized to the average intensity of individual experiments. For Figure 4D, by combining with actual observation, 0.15 of HsSAS6 fluorescence value (a.u.) was defined as a threshold to distinguish before and after HsSAS6 loading.

**Fig. S5.**
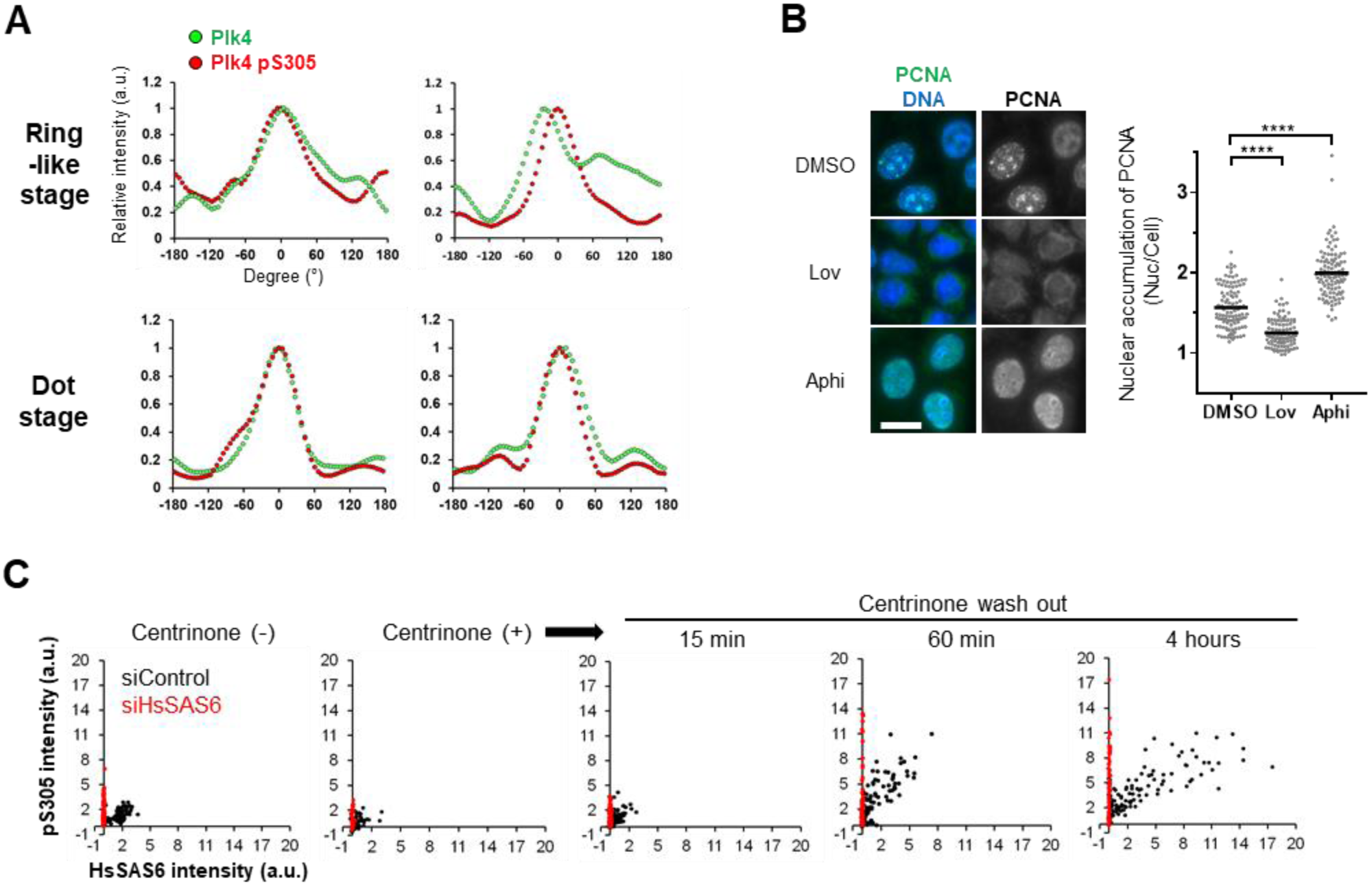
Analysis of Plk4pS305, Related to Fig. 4 and Fig. 5. **(A)** Relative intensities of Plk4 (Green) and pS305 (Red) in Figure 4B were quantified along Cep152 ring and maximum peaks of pS305 were set to 0°. Intensities were normalized to each maximum intensity. **(B)** Validation of cell cycle synchronization (Related to Figure 4E and 5) with chemicals by immunostaining against PCNA. Cells were treated with 6 μM aphidicolin (Aphi, 24 hours) and 10 μM Lovastatin (Lov, 20 hours). Nuclear accumulation of PCNA was calculated from the mean nuclear intensity divided by the mean cellular intensity. Compared to DMSO, Lovastatin suppressed nuclear accumulation of PCNA and aphidicolin increased nuclear PCNA in most cells, suggesting cell cycle synchronization in G1 and S phase, respectively. The graph shows mean value (black bar) of >100 cells. \*\**p* < 0.01, \*\*\*\**p* < 0.0001, ns, not significant, one-way ANOVA **(C)** Correlation of HsSAS6 and Plk4pS305 intensity. Data were the same as Figure 5D. Black and red dots represent the data from siControl and siHsSAS6-treated cells, respectively.

## Materials and Methods

### Plasmid construction

For recombinant protein expression in *E.coli*, cDNAs encoding human Plk4 full-length and a fragment (Kinase + L1: 1 - 585 a.a.) were cloned into pGEX 6p-1 and modified by inserting 6×His-tag cDNA sequence. The constructs were further modified by inserting cDNA sequence encoding mScarlet-I or GFP in between PreScission protease recognition site and Plk4 cDNA.

For over-expression of GFP-tagged Plk4 in HeLa cells, cDNAs encoding GFP and human Plk4 or Drosophila Plk4 were cloned into pcDNA5/FRT/TO. For weak expression of GFP-Plk4 in HeLa cells, the CMV2 promoter of pcDNA5/FRT/TO was replaced with human Plk4 promoter which was cloned from genomic DNA of HeLa cells, according to the previous study (*28*). For expression of GFP-Plk4-CAAX, pEGFPC1 (CMV promoter) was used and cDNA sequence encoding CAAX motif (derived from KRas4B) was inserted into the vector. 3×FLAG and Plk4-3×FLAG were expressed using pCMV15. Plk4 mutants such as kinase-dead (human: D154A, Drosophila: D156A), alanine mutants and phospho-mimetic mutants were generated by using PrimeSTAR mutagenesis basal kit (Takara).

### Protein purification

*E.coli* strain BL21 gold (DE3) was used for protein expression. Protein expression was induced at 18°C for 16 hours by incubating in LB medium supplemented with 0.3 mM IPTG. Cell pellets were suspended in lysis buffer (50 mM Tris (pH7.5), 300 mM NaCl, 2 mM MgCl2, 5 mM EDTA, 1 mM DTT, 0.5 mM PMSF, 0.5% TritonX-100) and lysed by lysozyme treatment and sonication. The lysates were then centrifuged at 9000 ×g for 45 min and the supernatants were collected. The supernatants were incubated with glutathione sepharose beads (GE healthcare) for 1 hour. The beads were washed with 1^st^ wash buffer (50 mM Tris (pH7.5), 800 mM NaCl, 2 mM MgCl2, 5 mM EDTA, 1 mM DTT, 0.5 mM PMSF, 0.5% TritonX-100) and then washed with pre-elution buffer (50 mM Tris (pH7.5), 500 mM NaCl, 5 mM β-mercaptoethanol). For GST-His6 or GST-Plk4(Kinase+L1)-His_6_ fragments, elution was performed in pre-elution buffer supplemented with 20 mM glutathione. For others, elution was performed by incubation with GST-tagged PreScission protease at 4°C, overnight. The eluates were incubated with Ni-Agarose beads (Wako) in binding buffer (50 mM Tris (pH7.5), 500 mM NaCl, 0.5% TritonX-100, 10 mM Imidazole, 5 mM β-mercaptoethanol) for 1 hour. The beads were washed with 2nd wash buffer (50 mM Tris (pH7.5), 500 mM NaCl, 50 mM Imidazole, 5 mM-βmercaptoethanol) and then proteins were eluted with elution buffer (50 mM Tris (pH7.5), 500 mM NaCl, 300 mM Imidazole, 5 mM β-mercaptoethanol). The eluates were dialyzed in dialysis buffer (20 mM Tris (pH7.5), 300 mM or 500 mM NaCl, 1 mM β-mercaptoethanol). Protein concentration was determined by Bradford assay.

### Spin-down assay

500 μM of GST-Plk4(Kinase+L1)-His_6_ fragments were incubated at 30°C for 1 hour in kinase buffer (20 mM Tris (pH7.5), 300 mM or 500 mM NaCl, 30 μM ATP, 5 mM MgCl_2_, 1 mM β-mercaptoethanol). The protein samples were incubated with GST-tagged PreScission protease at 4°C, overnight. After the reaction, the samples were centrifuged at 21500 ×g for 10 min and the supernatant was collected as “supernatant fraction”. The pellet was once washed with kinase buffer and then resuspended in kinase buffer (same volume as supernatant fraction) as “pellet fraction”. The fractions were analyzed by SDS-PAGE (10% polyacrylamide gel) and CBB staining using SimplyBlue SafeStain (ThermoFisher). Band intensities were measured using Fiji (NIH).

### Detection of protein aggregation

To detect the light scattering of protein solution, we used Prometheus NT.48 (NanoTemper technologies). 100 μg/ml of GST-His_6_ and GST-Plk4(Kinase+L1)-His_6_ fragments in buffer (20 mM Tris (pH7.5), 500 mM NaCl, 30 μM ATP, 2.5 mM MgCl_2_, 1 mM β-mercaptoethanol) were loaded into High sensitivity capillaries (NanoTemper technologies). According to the manufacturer’s instruction, the samples were subjected to a thermal ramp (15°C to 95°C, 1°C/min) with an excitation power of 100%. Data analysis was performed with the Prometheus PR. ThermControl software (NanoTemper technologies). Aggregation onset temperature was calculated from first derivative analysis on the software.

To detect the relative amount of protein aggregation, we used PROTEOSTAT protein aggregation assay kit (Enzo, ENZ-51023). 300 nM of protein samples in buffer (20 mM Tris (pH7.5), 300 mM NaCl, 1 mM β-mercaptoethanol) were supplemented with 0.1 mM MnCl_2_ and incubated at 37°C for 1 hour with or without lambda phosphatase (NEB, P0753S). According to the manufacture’s instruction, PROTEOSTAT detection reagent was added to each sample on 96-well clear bottom black plate (Corning, 3603). The fluorescence was detected using FilterMaxF5 (Molecular Devices) with excitation wavelength 535 nm and emission wavelength 595 nm.

### Human cell culture, cell synchronization and transfection

HeLa cells were obtained from the ECACC. Cells were cultured in DMEM containing 10% FBS and 1% penicillin/streptomycin at 37°C in 5% CO_2_ atmosphere. Cells were synchronized in G1 phase by incubating with 10 μM Lovastatin for 20 hours. For synchronization in S phase, cells were treated with 6 μM aphidicolin for 24 hours.

Transfection of plasmid DNA and siRNA was performed using Lipofectamine 2000 and Lipofectamine RNAiMAX (Life Technologies), respectively, according to the manufacturer’s instruction. Transfected cells were analyzed 48 hours after transfection with siRNA and 20, 24 or 48 hours after transfection with plasmid DNA.

### RNA interference

The following siRNAs were used: custom siRNA (Sigma Genosys) against 3’UTR of Plk4 (5’-CTCCTTTCAGACATATAAG-3’); Stealth siRNA (Life Technologies) against 3’UTR of HsSAS-6 (5’-GAGCUGUUAAAGACUGGAUACUUUA-3’); Silencer select Negative Control No.1 siRNA (Ambion, 4390843) and Stealth siRNA negative control Low GC Duplex #2 (Invitrogen, 12935110).

### Antibodies

The following primary antibodies were used; Rabbit polyclonal antibodies against Cep152 (Bethyl laboratories, A302–480A, IF 1:1000), STIL (Abcam, ab89314, IF 1:500), GFP (MBL, 598, IF 1:1000), Plk4 phospho-S305 (IF 1:500, WB 1:2000, the details are described below.) Mouse monoclonal antibodies against Plk4 (Merck Millipore, clone 6H5, MABC544, IF 1:500), HsSAS6 (Santa cruz Bio-technology, Inc., sc-81431, IF 1:500), Centrin-2 (Merck Millipore, clone 20H5, 04-1624, IF 1:1000), PCNA (Santa cruz Bio-technology, Inc., sc-56, IF 1:1000), GFP (Invitrogen, A11120, IF 1:1000), GST (MBL, M071-3, WB 1:2000) The following secondary antibodies were used; Alexa Fluor 488 goat anti-mouse IgG (H+L) (Molecular probes, A11001, 1:1000), Alexa Fluor 488 goat anti-rabbit IgG (H+L) (Molecular probes, A11008, 1:1000), Alexa Fluor 594 goat anti-mouse IgG (H+L) (Molecular probes, A11005, 1:1000), Alexa Fluor 568 goat anti-rabbit IgG (H+L) (Molecular probes, A11011, 1:1000), Goat polyclonal antibodies-HRP against mouse IgG (Promega, W402B, 1:5000), rabbit IgG (Promega, W401B, 1:5000) for WB.

Alexa 647-labeled anti-Cep152 (Bethyl laboratories, A302–480A) was generated using Alexa Fluor 647 antibody labeling kit (Molecular probes, A20186) and used for triple staining.

The Plk4pS305 antibody was generated by immunizing a rabbit with a S305-phosphorylated peptide corresponding to amino acids 301-314 (SISGpSLFDKRRLLC) of Plk4 (Eurofins Genomics). The antiserum was subjected to affinity-purification using the phosphorylated peptide and then absorbed non-specific antibodies using non-phosphorylated peptide.

### Chemicals

The following chemicals were used; Centrinone (MedChem Express, HY-18682), MG132 (Wako, 135-18453), MLN4924 (Chemscene, CS-0348), Aphidicolin (Sigma, A0781), Lovastatin (Merck, 438185).

### Imaging of fluorescence tagged proteins

Fluorescence-labeled protein samples in buffer containing 500 mM NaCl were mounted onto slide glasses (Matsunami, S0318) and covered with cover glasses (Matsunami, C015001). Images of aggregates were taken using Leica TCS SP8 inverted confocal microscope equipped with a Leica HCX PL APO × 63/1.4 oil CS2 objectives and excitation wavelength 488 nm or 561 nm.

### Immunostaining and imaging

HeLa cells cultured on coverslips (Matsunami, C015001) were fixed with cold Methanol at −20 °C for 7 min. The cells were washed with PBS for 5 min three times and incubated in blocking buffer (1%BSA, 0.05% Triton X-100 in PBS) for 30 min. The cells were then incubated with primary antibodies in blocking buffer at 4 °C, overnight and washed with PBS three times, and incubated with secondary antibodies for 1 hours at RT. The cells were washed with PBS twice and stained with Hoechst 33258 (DOJINDO) in PBS for 5 min at RT, and then mounted onto slide glasses.

For imaging of Figure 2C, S2, S3D and S4B, Leica TCS SP8 inverted confocal microscope equipped with a Leica HCX PL APO 63×/NA 1.4 oil CS2 objective was used. The images were collected at 0.4 μm z-steps. For Figure 3B, 3E, 4D, 5B-E, S3B, S4C-E, S5C, DeltaVision personal DV-SoftWorx system (Applied Prescision) equipped with a Olympus 60×/NA 1.42 oil objective and a CoolSNAP ES2 CCD camera or EDGE/sCMOS 5.5 camera was used. The images were collected at 0.2 μm z-steps. For Figure S5B, Zeiss Axio Imager M2 equipped with a 63×/NA 1.4 Plan-APOCHROMAT oil objective and a AxioCam HRm camera was used. The images were collected at 0.25 μm z-steps.

3D-SIM analysis in Figure 4 was performed using Nikon N-SIM imaging system equipped with Piezo stage, Apo TIRF 100×/NA 1.49 oil objective, iXon DU-897 EMCCD camera (Andor technology Ltd.) and excitation wave length 488 nm, 561 nm and 640 nm. The images were collected at 0.12 μm z-steps. Reconstitution of SIM images was performed using NIS-Element AR software.

Z-projection was performed using Maximum intensity projection using Fiji. The images in Figure 2C, 3B, 3E, 4B-E 5B, 5C, S2B, S3D, S4B, S4C and S5B show the maximum intensity projection and others are a single section.

### Quantification analysis

For counting *in vitro* aggregates per image (18.49×18.49 μm), fluorescence signals above the defined threshold intensity and size were regarded as aggregates and the number and area (size) were measured using Particle analysis in Fiji (NIH). For counting aggregates of GFP-Plk4(ΔCPB)-CAAX in HeLa cells, images were obtained by z-projection of a plasma membrane region (basal side, 0.4 μm Z-steps × 3). Then, fluorescence signals 3-fold higher than the mean intensity of the plasma membrane region were regarded as aggregates and the number was counted using Particle analysis in Fiji.

Fluorescence intensities of centriolar proteins were measured using Fiji by subtracting mean cytoplasmic signal intensity from mean centriolar signal intensity. A defined size of ROI was set for each experiments.

To quantify fluorescence patterns along mother centriole wall in SIM images, “oval profile” analysis in Image J (NIH) was used. Each maximus peak of Plk4pS305 intensity was set to the point of 0°. Intensities were normalized to each maximum intensity.

### Fluorescence Recovery After Photobleaching (FRAP)

For FRAP analysis, HeLa cells were cultured on 35 mm glass-bottom dishes (Greiner-bio-one, #627870). Before FRAP analysis, the culture medium was changed to DMEM containing 10%FBS without phenol red (Sigma).

FRAP analysis was performed using Leica TCS SP8 inverted confocal microscope equipped with a Leica HCX PL APO × 63/1.4 oil CS2 objectives and a 488 nm laser in a chamber in 5% CO2 at 37°C. The pinhole was adjusted at 2.0 airy units. Single section images were recorded at 1.29 sec (pre-bleach) and 5 sec (Fig S3F) or 10 sec (Fig 3A, D) (post-bleach) intervals. A region of interest (encircled with 2.54 μm diameter) around the centrosomal GFP signals was bleached with maximum laser power. Mean intensity values of the centrosomal GFP signals (encircled with 1.66 μm diameter) were measured using Fiji (NIH) and extracellular signals were subtracted as background. Signal intensity was normalized with the average of three pre-bleach signals. Due to centrosome movements, focus was readjusted during imaging.

### Immunoblotting

Protein samples were subjected to SDS-PAGE with 8% polyacrylamide gel and transferred onto Immobilon-P membrane (Millipore). The membrane was incubated in 5% skim milk in PBS and probed with the primary antibodies in 5% BSA in PBS, followed by incubation with their respective HRP-conjugated secondary antibodies. The membrane was washed in PBS containing 0.02% Tween. The signal was detected using Amersham ECL Prime Western blotting detection reagent (GE Healthcare) and a Chemi Doc XRC+ (BioRad).

### Protein sequence analysis

Intrinsically disordered regions and low-complexity regions were predicted using PrDOS and SMART respectively.

### Statistics

Statistical analyses were performed with GraphPad Prism 7.

